# Queuing models of gene expression: Analytical distributions and beyond

**DOI:** 10.1101/2020.03.04.976738

**Authors:** C. Shi, Y. Jiang, T. Zhou

## Abstract

Activation of a gene is a multistep biochemical process, involving recruitments of transcription factors and histone kinases as well as modification of histones. Many of these intermediate reaction steps would have been unspecified by experiments. Therefore, classical two-state models of gene expression established based on the memoryless (or Markovian) assumption would not well describe the reality in gene expression. In fact, recent experimental data have indicated that the inactive phases of gene promoters are differently distributed, showing strong memory. Here, we use a non-exponential waiting-time distribution to model the complex activation process of a gene, and analyze a queuing model of stochastic transcription. We successfully derive the analytical expression for the mRNA distribution, which provides insight into the effect of molecular memory created by complex activating events on the mRNA expression. We find that the reduction in the waiting-time noise may result in the increase in the mRNA noise, contrary to the previous conclusion. Based on the derived distribution, we also develop a method to infer the waiting-time distribution from a known mRNA distribution. Data analysis on a realistic example verifies the validity of this method.

**SIGNIFICANCE:** Activation of a gene is a complex biochemical process and involve several intermediate reaction steps, many of which have been unspecified by experiments. Stochastic models of gene expression that were previously established based on the constant reaction rates would not well reflect the reality in gene expression. To this end, we study a queuing model of stochastic transcription which assume that the reaction waiting time follows a general distribution and derive the analytical expression for mRNA distribution. Our results provide insight into the role of molecular memory in fine-tuning the gene expression noise, and can be used to infer the underlying molecular mechanism.

## INTRODUCTION

Gene expression is a complex biochemical process, inevitably leading to stochastic fluctuations in mRNA and further protein [1-4]. While this inherent noise would be important for the maintenance of cellular functioning and the generation of cell phenotypic variability, mathematical models are a strong tool to quantify the contributions of different noisy sources of gene expression.

Single-cell studies on gene expression have indicated that most genes in prokaryotic and eukaryotic cells are expressed in a bursty fashion [5,6]. In theory, this kind of expression manner may be modeled by the two-state model where the gene switches between two states: one transcriptionally active (on) state and one transcriptionally inactive (off) state. Classical two-state models of gene expression [7-20] assume that all the involved reaction rates are constants, implying that the reaction waiting times, particularly including those from the on to off states and vice versa, follow exponential distributions. This Markovian assumption has led to important successes in the modeling of many gene expression processes [21,22]. With the Markovian assumption, analytical mRNA or protein distributions have also been derived [7,10,11,23-25], which provide the best description of the stochastic properties of the gene systems. These studies have revealed that the characteristic parameters in the reaction waiting-time distributions can have a significant impact on the mRNA and protein levels.

However, biochemical events involved in gene expression occur not necessarily in a Markovian manner but may take place in a non-Markovian manner. First, the complex control process of transcription initiation can generate non-exponential time intervals between transcription windows [26-28]. Second, the synthesis of an mRNA would involve multiple intermediate reaction steps that have been unspecified due to experimental technologies, creating a memory between individual reaction events [30,31]. For example, the inactive phases of promoter involving the prolactin gene in a mammalian cell are differently distributed, showing strong memory [26]. Indeed, the increasing availability of time-resolved data on different kinds of interactions has verified the extensive existence of molecular memory in biological systems [26-28, 32-34]. How non-Markovian reaction kinetics impact gene expression remains elusive even though non-Markovian models of gene expression have been established in terms of chemical master equations (CMEs) [20,35].

There have been studies on non-Markovian models of stochastic gene expression, which can be divided into two classes from the viewpoint of continuous time random walk (CTRW) [35,36]: active CTRWs where waiting time needs to be reset, and passive CTRWs where waiting time cannot be reset. Qiu, et al. [37], studied a specific active CTRW model of gene expression and derived the analytical expression for stationary gene-product distribution. In contrast, more studies focused on queuing models of gene expression (belonging to the passive CTRW class). Pedraza and Paulsson [31] studied a queuing model of gene expression from the viewpoint of queue theory, showing that common types of molecular mechanisms can produce gestation and senescence periods that reduce noise without requiring higher abundances, shorter lifetimes, or any concentration-dependent control loops. Kulkarni, et al. [38,39] analyzed a stochastic model of bursty gene expression which considers general waiting-time distributions governing arrival and decay of proteins. By mapping the gene system to standard queuing models, they also derived analytical expressions for the steady-state protein noise (defined as the ratio of the variance over the squared mean). In addition, previous studies [31,40] showed that a reduction in the noise of the waiting time distribution of the inactive state leads to a reduction in the noise of mRNA distribution. Nevertheless, for queuing models of gene expression, analytical mRNA or protein distributions have not been derived by far.

In this paper, we also study a queuing model of stochastic gene expression, focusing on gene-product distribution rather than on statistical quantities such as the mean and variance studied in ref. [26,39-41]. We derive the analytical expression for stationary gene-product distributions, which provide insight into the role of molecular memory in fine-tuning the mRNA or protein noise and properties of the steady states. We also present estimations on the bounds of the expression noise constrained by the mean and variance of the waiting-time distribution. In addition, we develop an effective method to infer a waiting-time distribution and further the promoter structure from a known mRNA or protein distribution obtained by experimental methods such as mRNA-FISH labeling or RT-qPCR [42,43]. Our inference method is different from previous inference methods [27,40,42-45], which used the qualitative properties of the gene expression noise to infer microscopic kinetic parameters, such as transcription and translation rates. Data analysis on a realistic example verifies the validity of this method.

## MATERIALS AND METHODS

The common two-state model of stochastic transcription assumes that the gene promoter has two activity states: one active (on) state in which the gene is transcribed and the other inactive (off) state in which transcription is prohibited. These transcriptional active and inactive states may switch to each other. The mRNA molecules transcribed from DNA degrade in a linear manner. All involved reaction rates are assumed as constants, implying that the reaction waiting times follow exponential distributions or the reaction events take place in a Markovian (memoryless) manner [21,22]. Different from the classical on-off model, our model assumes that the waiting time from the off state to the on state follows a general distribution denoted by *f*(*t*). This assumption is reasonable since the turning on of transcription is a complex process with intermediate reaction steps that would have been unspecified by experiments. We point out that although this model has been studied in terms of statistical quantities [26,39,40], the analytical mRNA distribution has not been derived by far.

For the modeling convenience, we list all biochemical reactions as follow

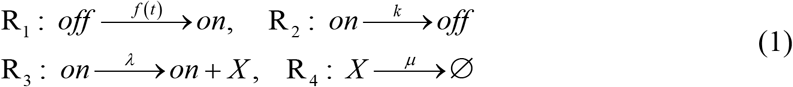

where *X* may represent mRNA or protein (without loss of generality, we assume that it represents mRNA), *k* is the switching rate from on to off states, *λ* represents the transcription rate, and *μ* represents the degradation rate of mRNA. These reaction rates are assumed to be constants, implying that the corresponding reaction waiting-time distributions are exponential.

Let *S*(*t*) record the elapsed time since the gene enters the off state at time *t*, and let *U*(*t*) denote the promoter state at time *t*. Then, the state of the system at time *t* can be described by the following time-evolutional probabilities

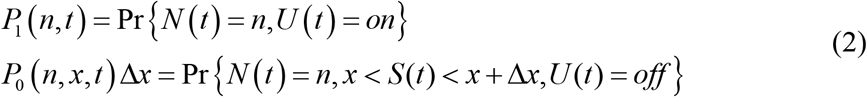

where *x* represents the elapsed time, *P*_0_(·) and *P*_1_(·) represent the probability density functions in off and on states respectively. In order to derive the renewal equations for *P*(*n, t* + Δ*t*) and *p* (*n, x, t* + Δ*t*) with regard to time increment Δ*t*, we first interpret two conditional probabilities [41,46,47]: *H*(*x* + Δ*t*)*H*(*x*) and *f*(*x*)ļ*H*(*x*), where 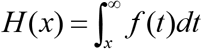 is the survival function of off state. The former represents the conditional probability that reaction R_1_ in Eq.(1) does not occur within time interval (*x,x*+Δ*t*) if the elapsed time is *x*, whereas the latter represents the instantaneous probability that reaction R_1_ happens also if the elapsed time is *x*.

According to the total probability principle, we can establish the following renewal equation for *P*_1_(*n,t*) and *P*_0_(*n,t*):

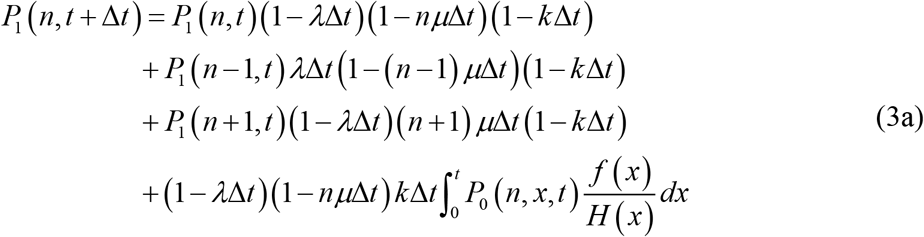

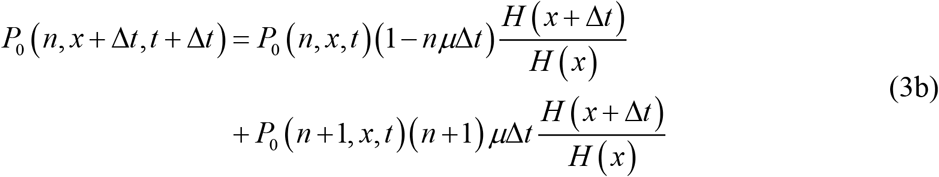

In Eqs. (3a) and (3b), *P_i_*(·) denotes the probability set Pr{·} defined in Eq. (2). The following boundary condition at *x* = 0 needs to be imposed due to consideration of the probability equilibrium

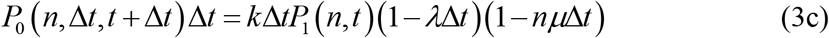

Note that in the limit of small Δ*t*, Eqs. (3a) and (3b) will become differential equations. If the stationary distributions of *P*_1_(*n,t*) and *P*_0_(*n,x,t*) exist, and are denoted by *P*_1_(*n*) and *P*_0_(*n,x*) respectively, Eqs. (3a) and (3b) at steady state become

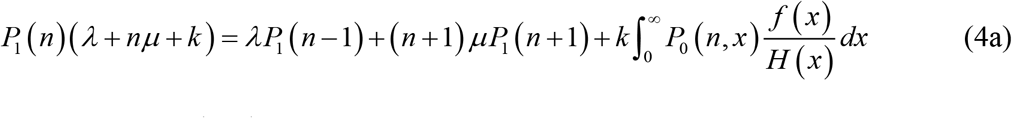

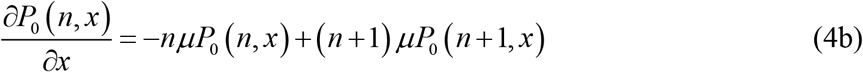

whereas Eq. (3c) becomes

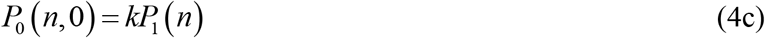

In order to solve Eqs. (4a) and (4b) with Eq. (4c), we introduce the probability generating functions as done previously

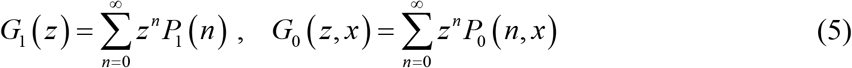

and denote *W*_0_ (*z, x*) = *G*_0_(*z, x*)/*H*(*x*). Then, we can obtain the following differential equations

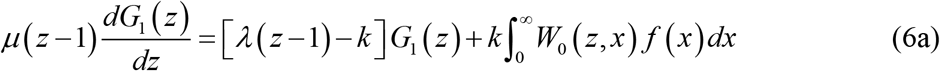

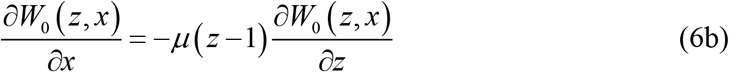

with the boundary condition

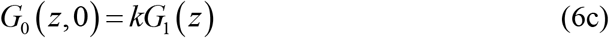

One will see that Eqs. (6a)-(6c) are analytically solvable. Thus, we can obtain the total probability-generating function *G*(*z*) = *G*(*z*) + *G*_0_(*z*) with 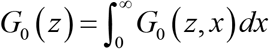 and further the total stationary probability distribution *P*(*m*) = *P*_0_(*m*) + *P*_1_(*m*).

## RESULTS

### Analytical distributions

Using the method of characteristic line, we can derive the solution of *W*_0_(*z, x*) from Eqs. (6b) and (6c)

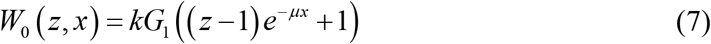

If we denote 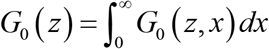, then Eq. (7) becomes

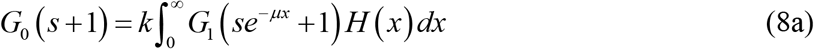

and Eq. (6a) becomes

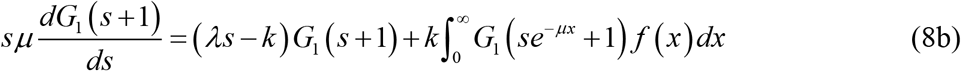

where *s* = *z*−1. By making Taylor expansions 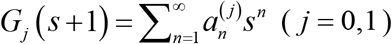, making use of the binomial method [48,49], and noting the total probability-generating function *G*(*s*)=*G*_0_(*S*)+*G*_1_(*s*) and the total probability *P*(*m*)=*P*_0_(*m*)+*P*_1_(*m*) as well as the relationship between the probability-generating function and the probability distribution, i.e., *P*(*m*) = (1/*m*!)[*d^m^G*(*s*)/*ds^m^*]|_*s*=−1_, we can finally reach the following formal expression for the stationary mRNA distribution (see Appendix A for derivation)

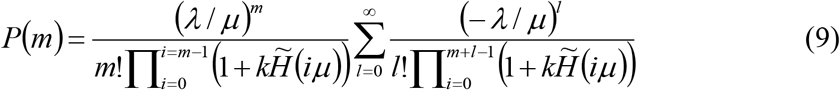

where 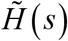 represents the Laplace transform of *H*(*t*), i.e., 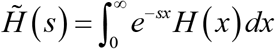.

In particular, if the waiting time from off to on state follows an exponential distribution of the form *f*(*t*) = *re^−rt^*, then 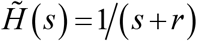. In this case, we can obtain from Eq. (9) the following analytical mRNA distribution

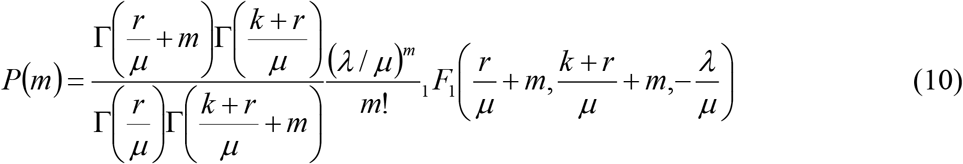

where _1_*F*_1_ denotes the confluent hypergeometric function [50]. This distribution was ever derived in previous works [10,51,52].

Next, we use Eq. (9) to calculate the mRNA distribution for a two-state model with an arbitrary inactive pathway (referring to Fig.1A). The probability density function for the waiting time takes the form *f*(*t*) = **a**_0_ exp(**A***t*)**u**_0_ (seeing Appendix B or referring to [10,27,40]), where **A** (a matrix) describes transitions among off states, **u**_0_ (a column vector) describes the transitions from the off states to the on state, and **a**_0_ (a row vector) denotes the initial condition. The Laplace transform of probability density function *f*(*t*) is given by 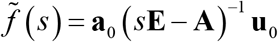, and the Laplace transform of survival probability function *H*(*t*) is given by

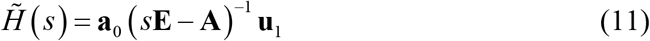

where **u**_1_ =(1,1,⋯,1)^T^ is a column vector. Substituting Eq. (11) into Eq. (9), we obtain the mRNA distribution for the two-state model with an arbitrary inactive pathway.

**Fig. 1.**
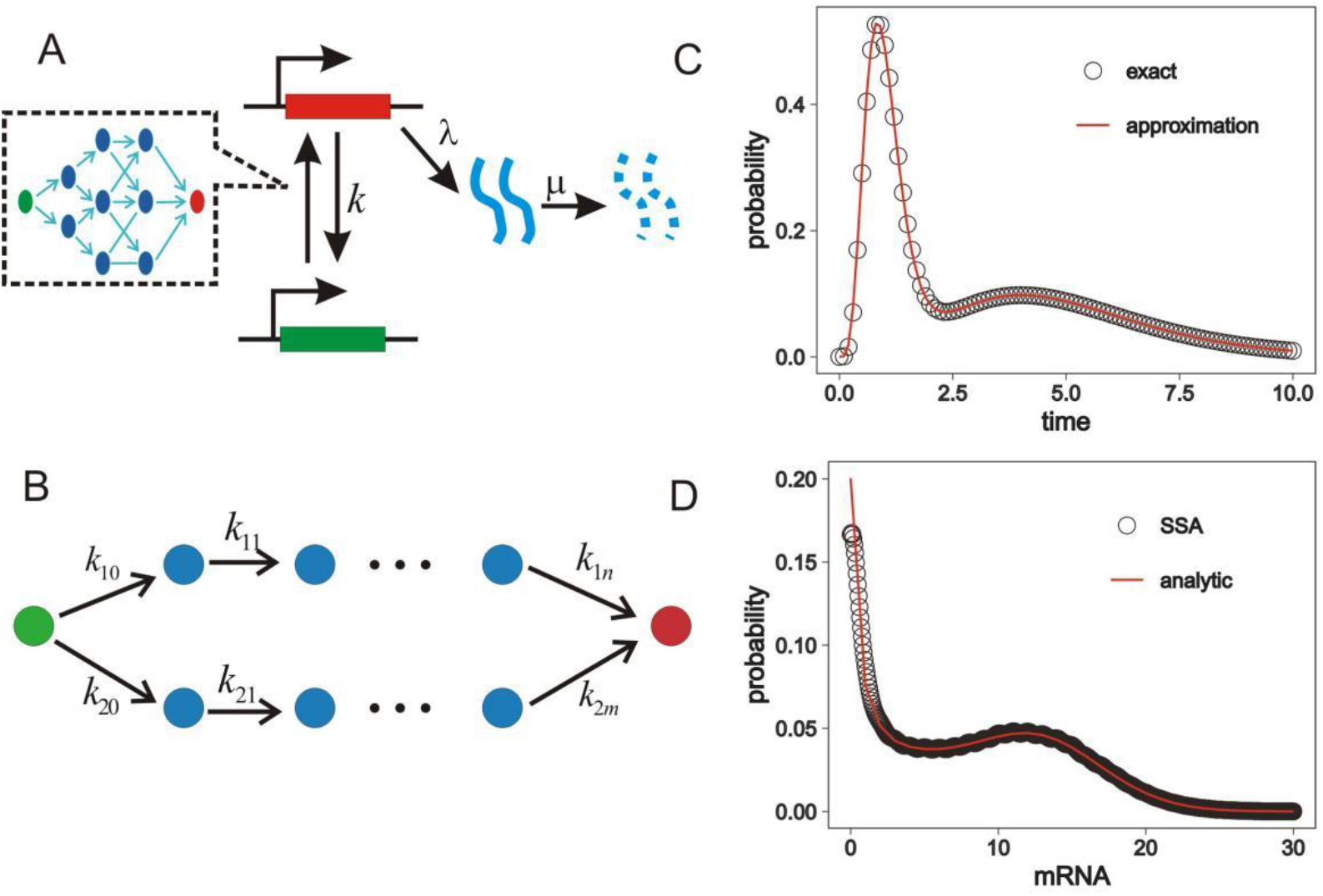
(A) A two-state model of gene expression with an arbitrary inactive pathway; (B) Two parallel inactive pathways; (C) Waiting-time distribution, where red line represents an approximate solution obtained using a mixture Gamma distribution, empty circles represent the exact solution obtained by the Gillespie algorithm, the parameter values are set as *k*_10_ = *k*_20_ = 50, *k*_11_ = *k*_12_ =…= *k*_15_ = 1, *k*_21_ = *k*_22_ =…= *k*_26_ = 6; (D) mRNA distribution, where red line represents the analytical solution,empty circles represent the numeric solution obtained by the Gillespie algorithm, parameter values are set as *k* = 1/3, *λ* = 15, *μ* = 1, and the other parameters are same as (C).

When the inactive pathway consists of two parallel pathways (referring to Fig. 1B), the Laplace transform of waiting time distribution is given by

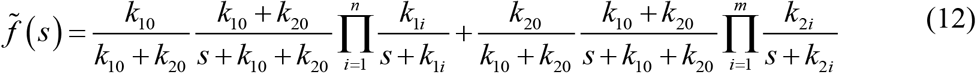

Under the assumption of *k*_11_ = *k*_12_ =⋯= *k*_1*n*_ ≡ *k*_1_, *k*_21_ = *k*_22_ =⋯= *k*_2*m*_ ≡ *k*_2_, and *k*_10_ + *K*_20_ >> max {*k*_1_, *K*_2_}, we find that the waiting-time distribution can be approximated by a mixture Gamma distribution of the form (referring to Fig. 1C)

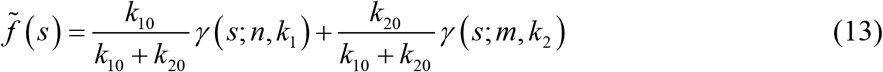

where 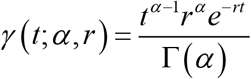 is a common Gamma distribution. The valid of this approximation has numerically been verified, referring to Fig. 1D where empty circles represents the exact mRNA distribution obtained by the Gillespie algorithm [53] whereas the line represents the approximate mRNA distribution obtained using Eq. (13).

### Effect of molecular memory on the mRNA noise

Using the derived-above analytical mRNA distribution for the gene model with a general waiting-time distribution, we find that the mRNA noise takes the following form

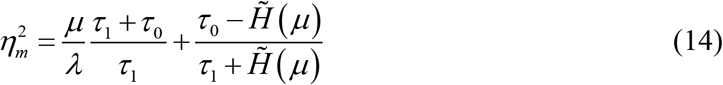

where *τ*_1_ = 1/*k* represents the average waiting time in the on state whereas 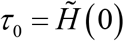 represents the average waiting time in the off state. For the gene model above, this formula is exact since the corresponding mRNA distribution is exact. Moreover, the mean mRNA is given by 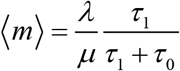. If we consider that the waiting time in the off state follows a Gamma distribution *γ*(*t;γ,r*), we have 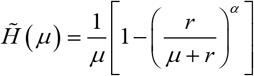. When *α* = 1, the model reduces to the case of exponential distribution and the corresponding formula for the mRNA noise becomes

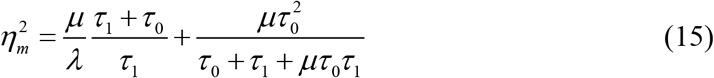

which is a known result [29,40,51,54].

Under the assumption that the waiting time follows a Gamma distribution, Eq. (14) can be rewritten as

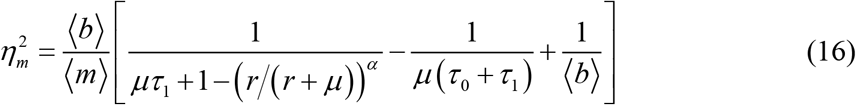

If the degradation of mRNA is slower or the waiting time in the on state is shorter (*μτ*_1_ <<1), the above calculation formula can be approximated as

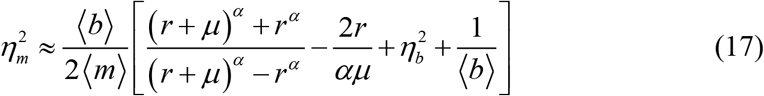

where 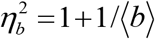 represents the noise of burst, 〈*b*〉 = *λ/k* represents the mean of burst size. Formula (17) is same as the result derived by Schwabe et al [40], which can reduce to the formula derived by Pedraza and Paulsson [31] in the absence of degradation.

Previous studies showed that if the waiting time follows a Gamma distribution, a reduction in the noise of the waiting time distribution in the off state leads to a reduction in the noise of mRNA distribution [15,31,40]. If considering another kind of waiting-time distribution, then we obtain a different result. As seen in the above, the waiting-time distribution can be well approximated by a mixture gamma distribution if the inactive process consists of two parallel pathways. Now, we assume that the waiting time follows a mixture gamma distribution given by

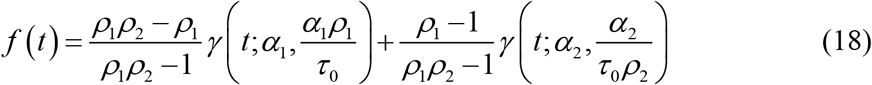

where *ρ*_1_ > 1 and *ρ*_2_ > 1 are constants. Then, it is not difficult to show that the mean of the waiting time is 〈*T*〉 = *τ*_0_, and the variance of the waiting time is

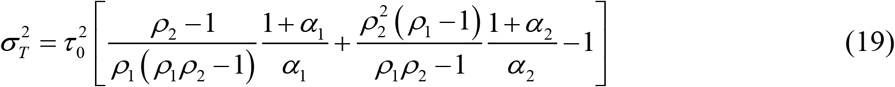

Changing the parameter *ρ*_1_ and *ρ*_2_ allow us tune the noise of waiting-time distribution 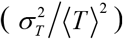 while keeping the mean of waiting time fixed. For clarity, we set *ρ*_1_ = (*c* + 1)/*c* and *ρ*_2_ = (2−*c*)/(1−*c*) with 0 < *c* < 1. With this setting, we find that with the increase of parameter *c*, the noise of the waiting-time distribution increases (Fig. 2A) but the mRNA noise decreases (referring to Fig. 2B). Alternatively, a reduction in the noise of the waiting time leads to an increase in the mRNA noise (Fig. 2C), which is contrary to the previous conclusion. The possible reason is that the mRNA noise can vary in a wide range even if the mean and noise of the waiting time are fixed (Fig. 2C).

**Fig. 2.**
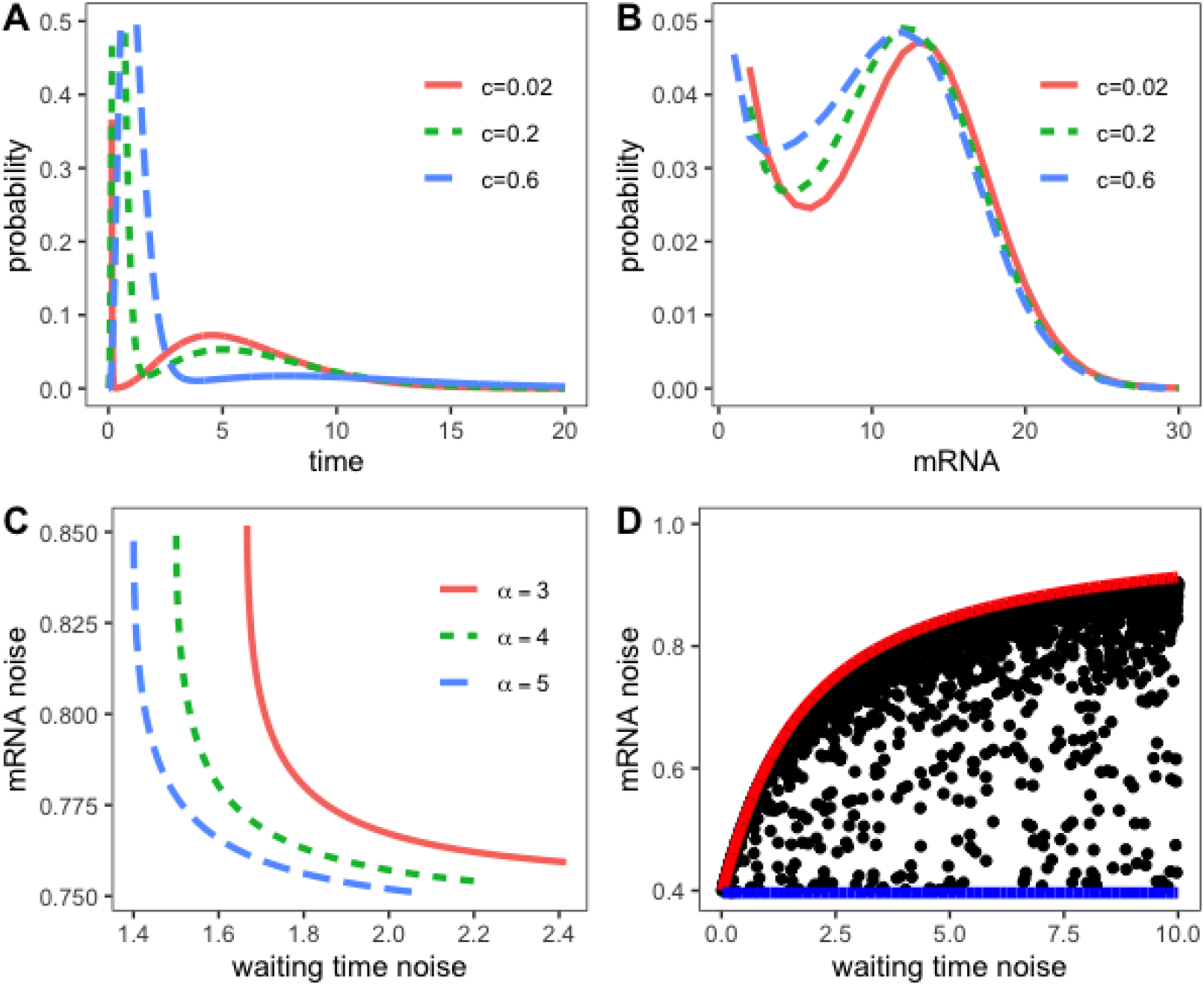
The mRNA noise adjusted by molecular memory. (A) Waiting-time distributions for different values of *c*, where *τ*_0_ = 3, *k* = 1/3, *α*_1_ = *α*_2_ = 4, *λ* = 15, *μ* = 1; (B) mRNA distribution for different values of *c*, where the parameter values are set as the same as (A). Here the probability of the zero mRNA is not shown; (C) mRNA noise versus waiting-time noise for different values of *c*, where *α*_1_ = *α*_2_ = *α* = 3,4,5 are set, and other parameter values are the same as (A); (D) mRNA noise versus waiting-time noise, where *b*_1_,*b*_2_,*α*_1_,*α*_2_ are uniformly sampled from the interval (1,200), parameter values are set as *τ*_0_ = 2, *k* = 0.5, *λ* = 15, *μ* = 1, the red and blue lines respectively represent the upper bound and lower bounds of the mRNA noise calculated using Eq. (21).

Next, we estimate the bounds of the mRNA noise in the case that the mean and noise of waiting time are fixed. For this, we first need to estimate the bounds of 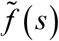. By calculation, we can obtain the following estimation (seeing Ref. [55])

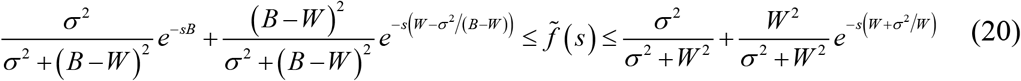

where *B* is the upper bound of the waiting time *T, W* and *σ*^2^ are the mean and variance of the waiting time, respectively. The upper bound corresponds to a two-point mixture distribution with mass *σ*^2^/(*σ*^2^ + *W*^2^) on position 0 and mass *W*^2^/(*σ*^2^ + *W*^2^) on position *W*+*σ*^2^/*W*. The lower bound corresponds to a two-point mixture distribution with mass *σ*^2^/(*σ*^2^+*W*^2^) on position 0 and mass *W*^2^/(*ρ*^2^+*W*^2^) on position *W+σ*^2^/(*σ*^2^(*B−W*)^2^) on position *B* and mass (*B−W*)^2^/(*σ*^2^(*B−W*)^2^) on position *W−σ*^2^/(*B−W*). When *B*→∞, the estimation becomes

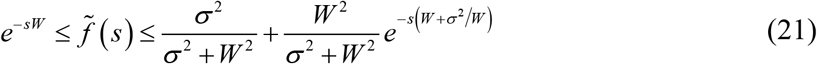

After having had the estimation of 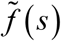, we can further obtain the estimation for the bounds of first 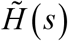 and then the mRNA noise. To numerically verify the resulting estimations for the bounds of the mRNA noise, we randomly sample parameters (*ρ*_1_,*ρ*_2_,*α*_1_,*α*_2_) while keeping the other parameters in the mixture distribution (referring to Fig. 2D) fixed. From this figure, we observe that with the increase in the waiting-time the noise, the lower bound of the mRNA noise is almost a constant but the upper bound increases.

### Inferring waiting-time distribution from mRNA distribution

The mRNA distribution can be obtained through experimental techniques such as mRNA-FISH labeling or RT-qPCR on individual cells. An interesting question is whether we can infer the underlying transcription mechanism from experimental data. In this section, we will provide a method to infer the waiting-time distribution from the known mRNA distribution. Without loss of generality, we set the mRNA degradation rate to be unit.

We first introduce binomial moments of the stationary mRNA distribution *P*(*n*), which are defined as [48,49]

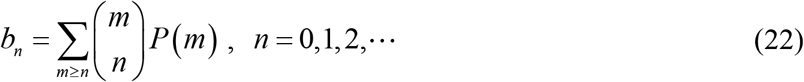

where *b*_0_ = 1 due to the probability conservation, and symbol 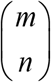 represents the combinatorial number of arbitrarily choosing *n* numbers from *m* numbers. Based on these binomial moments *b_n_*, we further introduce a new index (see Appendix C for the reason of introduction)

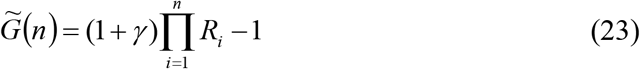

where 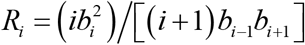, and *γ* is a constants larger than 0. Note that Distribution *P*(*n*) can be obtained from experimental data, so can binomial moments *b*(*n* = 0,1,⋯,*N*+1), all *R_i_* and all 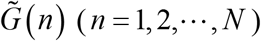. Furthermore, we can obtain first the continuous-variable function 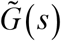 (representing the Laplace transform of the discrete-variable *G*(*n*)) using the Burlisch-Stoer rational interpolation in the R package “pracma” [56], and then *G*(*t*) by the numerical inverse Laplace transform of 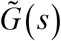 using the function “invlap” also in the R package “pracma”. For *t* ≥ *δ*_0_ (*δ*_0_ is a very small number), we can prove that *H*(*t*) = *c*_0_*G*(*t*), where *c*_0_ is a constant (≈ 1/*G*(*δ*_0_)) (seeing Appendix C for detail). Therefore, we can reconstruct an approximate waiting-time distribution according to *f*(*t*) = −*dH*(*t*)/*dt*. We point out the parameter *γ* does not influence the shape of the resulting distribution, so we can set it as a constant of more than 0. In addition, we can determine two parameter values: first 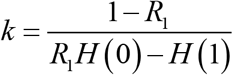 (the mean switching rate from on to off states) and then *λ*=*b*_1_(1+*kH*(0)) (the mean transcription rate). To that end, we finish our inference method. We emphasize that in contrast to the previous inference method, our inference method do not need to assume the type of waiting-time distribution in advance.

To verify the above inference method, we generate the mRNA distributions for the two-state model with a different waiting-time distribution given by Eq. (9). For gamma distribution with *α*=1 (exponential distribution), the predicted distribution and the exact distribution overlap even for a small *N* (Fig. 3A). For *α*> 1, the predicted distribution gradually tends to the exact distribution with the increase of *N* (Fig. 3B). The larger the *α* is, the larger *N* is required to infer a ‘better’ distribution. Actually, we can infer the exact distribution, provided that the mRNA distribution is precisely measured. However, the error of *b_k_* becomes larger with the increase of order *k* for any mRNA distribution generated from numerical simulation or obtained by experiments, so it is difficult to infer an exact distribution, particular in the case of strong memory. Although the shape of waiting-time distribution do not significantly affect the mRNA distribution, our result indicates that the information on waiting time can actually be encoded in an mRNA distribution.

**Figure 3.**
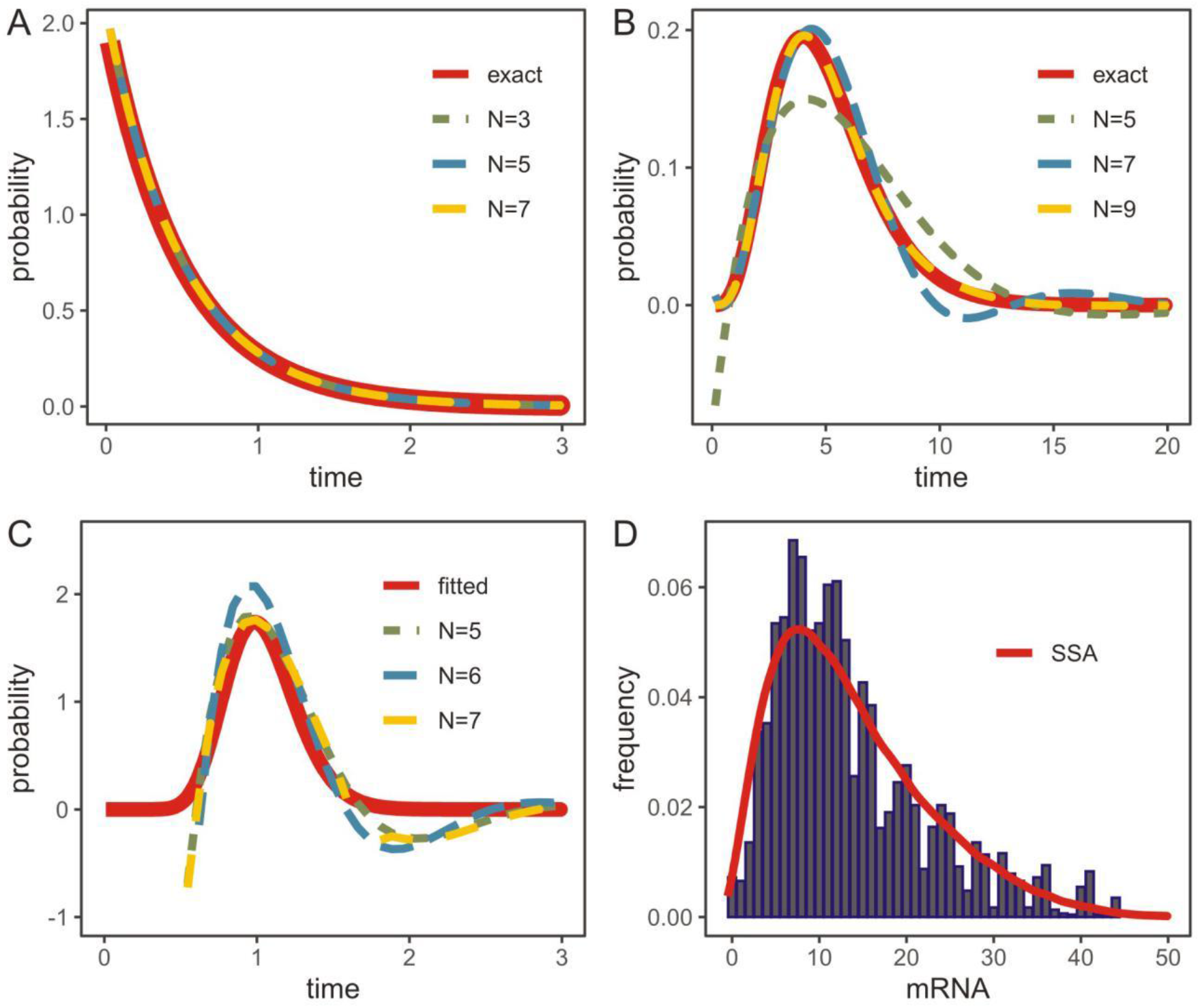
Inferring waiting-time distribution from mRNA distribution. (A) the parameter values corresponding to the exact distribution (red line) are as: *τ*_0_ =0.5, *k* =2, *α*=1, *λ*=5, *μ*=1; (B) the parameter values corresponding to the exact distribution (red line) are set as *τ*_0_ = 5, *k* = 2, *α*= 5, the other parameter values are the same as in (A); (C) the waiting-time distribution inferred from the experimental mRNA distribution for PDR5 gene, where the red line denotes the distribution fitted using Gamma distribution; (D) Comparison between observed data [42] (blue histogram) and the mRNA distribution obtained by stochastic simulation (red line), where the parameter values are set as *α*=18, *r* =17.75, *k* =2, *λ*=41.5, *μ*=1.

Finally, we apply the above inference method to a realistic example. We try to infer the waiting-time distribution from the mRNA distribution for gene PDR5 measured by Zenklusen et al [42]. The inferred mRNA distribution is demonstrated in Fig. 3C. Here, we fit the predicted waiting-time distribution (*N* = 5) with a Gamma distribution, and determine the left parameters of the model (Appendix C). The mRNA distribution obtained through the stochastic simulation well fits to the experimental data (Fig. 3D). We observe that there are errors between simulated and exact distributions, but if more binomial moments are used, then these errors will become smaller. In fact, we have the reconstruction formula: 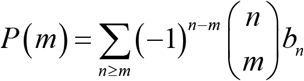 and binomial moments tend to zero as their orders go to infinity [48]. We point out that although the exact waiting-time distribution is difficult to infer, our numerical result indicates that the waiting-time distribution is definitely unimodal (referring to Fig. 3C). This qualitative result is independent of how many binomial moments are used to reconstruct the mRNA distribution.

## CONCLUSIONS AND DISCUSSION

In this paper, we have derived the analytical stationary mRNA distribution for the two-state model of gene expression where the waiting time from off to on states is assumed to follow a general distribution whereas the waiting time from on to off states is assumed to follow an exponential distribution. This derived distribution can provide insight into the role of molecular memory characterized by non-exponential waiting-time distribution in fine-tuning the gene expression noise. In contrast to previous methods [7,10,11,23,24], our method is not limited to a specific Markov chain description of various possible promoters containing transcriptionally active and inactive states. Actually, our derivation method can be applied to two-state gene models with arbitrarily many parallel or crosstalk inactive pathways.

Previous studies showed that a reduction in the waiting-time noise can lead to a reduction in the mRNA noise [31,40], under the assumption that the waiting time follows a Gamma distribution. In this paper, however, we found that a reduction in the waiting time noise may lead to the increase in the mRNA noise, under the assumption that the waiting time distribution follows a mixture Gamma distribution, which can model the case that the inactive process consists of two parallel inactive pathways. The plausible reason is that the mRNA noise may vary in a wide range even if the mean and noise of waiting time are fixed. For the two-state model of gene expression, we only considered that the waiting-time distribution for the off-to-on reaction is general. Actually, waiting-time distributions for on-to-off reaction, transcription reaction, and degradation reaction may also be non-exponential [29,31]. For these cases, how mRNA or protein distributions are derived as well as how molecular memory affects the expression level and noise needs further studies.

Previous results also showed that the observed mRNA distribution contains only the limited information on the transcriptional mechanism [40], and can be fitted by the two-state model with different parameters. In contrast, our results showed that the information on the waiting time is actually encoded in the mRNA distribution. In particular, we can infer the exact waiting-time distribution, provided that the mRNA distribution is precise. Our method can also be used to infer the waiting-time distribution from the mRNA distribution obtained by experimental data. For example, we have applied the method to the PDR5 gene [42] and found that the waiting-time distribution is unimodal. But this qualitative result expects experimental verification.

## Appendix A Derivation of analytical mRNA distribution

By expanding the probability-generating function 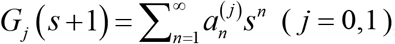, we can derive the following relations

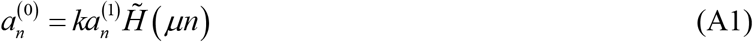

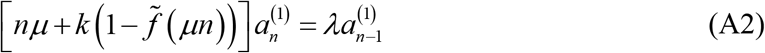

where 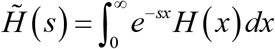 and 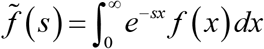. Note that 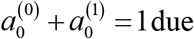 to the conservative condition of probability. We get the expression of 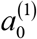 from Eq. (A1) as following:

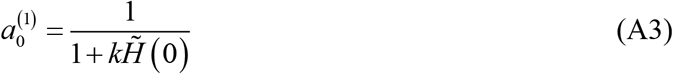

Since the total probability distribution of mRNA is *P*(*m*) = *P*_0_(*m*)+*P*_1_(*m*), the corresponding total binomial moments are 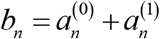. Combing Eqs. (A1)-(A3), we obtain the following expression of *b_n_*:

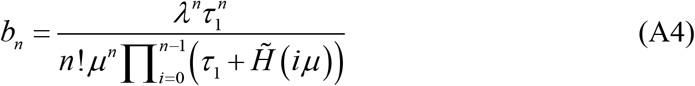

where *τ*_1_ = 1/*k* represent the average waiting time in the on state. With the expression of *b_n_* at hand, the corresponding distribution can be calculated according to the reconstructing formula 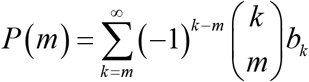 [48,49], where 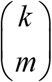 represents the combinatorial number of arbitrarily choosing *m* numbers from *k*, and *b_k_* represent the binomial moments. Thus, the stationary distribution of the mRNA *P*(*m*) is given by

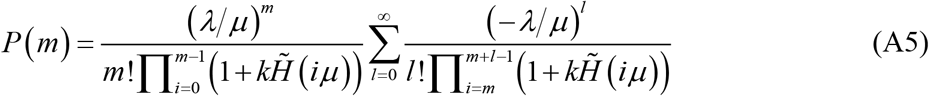

where *m* = 1,2,⋯.

## Appendix B Calculation of waiting-time distribution

For the inactive pathway shown in Fig. 1B, the probability distribution of waiting time is calculated according to *f*(*t*) = **a**_0_ exp(**A***t*)**u**_0_, where matrix **A** is

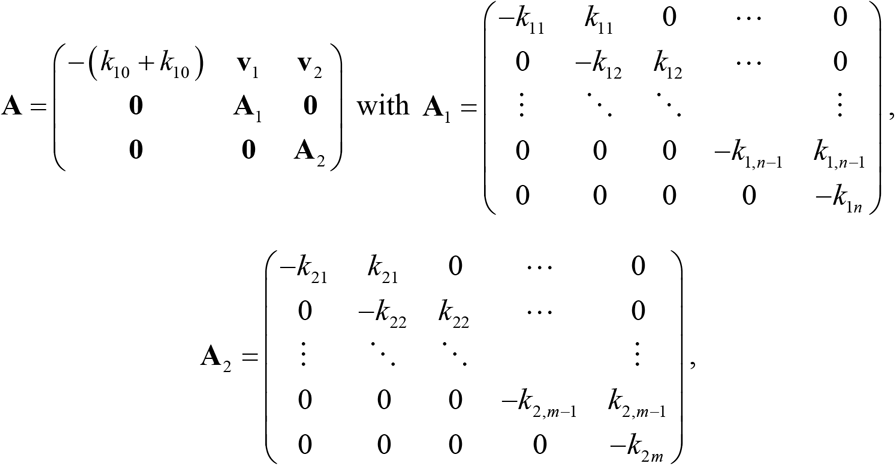

and **v**_1_ =(*k*_10_,0,⋯,0), **V**_2_ = (*k*_20_,0,⋯,0). The column vector **u**_0_ takes the form: **u**_0_ =(0,⋯,0,*k*_1*n*_,0,⋯,0,*k*_2*m*_)^T^, the row vector **a**_0_ takes the form: **a**_0_ = (1,0,⋯,0). According to 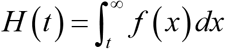, we thus have *H*(*t*) = **a**_0_ exp(**A***t*)**A**^−1^**u**. The Laplace transform of *f*(*t*) is calculated according to 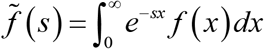. Thus, we have 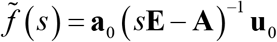, where **E** is the unit matrix. Similarly, we can have 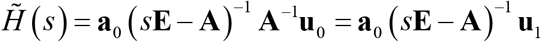, where **u**_1_ = (1,1,⋯,1)^T^.

## Appendix C A method to infer the waiting-time distribution from experimental data

Note that binomial moments *b_i_* (*i* = 1,2,⋯, *N*) can be estimated from experimental data. Thus, 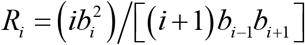 are known. Setting *μ* = 1 and substituting these expressions of *R_i_* into Eq. (A4), we obtain the following relationship

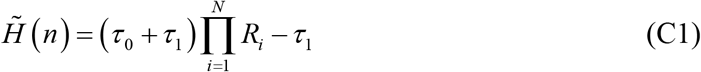

where *τ*_0_ and *τ*_1_ represent the mean waiting time of the OFF and ON states, which are experimentally measurable quantities and are therefore known. By smoothing discrete variable function 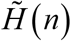 through interpolation, we can obtain continuous variable function 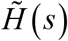. Furthermore, we can obtain survival function *H*(*t*) through the inverse Laplace transform.

If we set *τ*_0_ =*γτ*_1_(*γ* is a positive constant of more than) and *τ*_1_ = 1, then

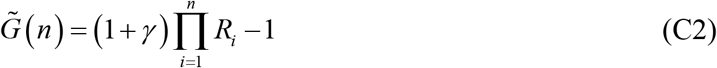

By smoothing discrete variable function 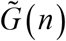 through interpolation, we can obtain continuous variable function 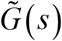. Furthermore, we can obtain the function *G*(*t*) through the inverse Laplace transform.

Combining Eq. C1 and Eq.C2, we obtain the following relation:

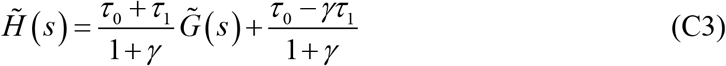

Applying inverse Laplace transform to Eq.C3, we have:

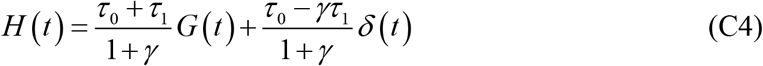

where *δ*(*t*) denotes the Dirac delta function. Therefore, for *t* ≥ *δ*_0_ (*δ*_0_ is a very small number), we obtain *H*(*t*) = *c*_0_*G*(*t*). Since *H*(*δ*_0_) ≈ 1, we have *c*_0_ ≈ 1/*G*(*δ*_0_).

Once the survival function *H*(*t*) is obtained in such a manner, we can further determine the values of other parameters in the model. From Eq. (C1), we can determine the value of *k* as 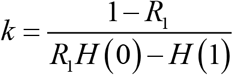. Thus, the value of *λ* can be determined according to *λ*=*b*_1_(1+*kH*(0)).

## AUTHOR CONTRIBUTIONS

C.S., Y.J. and T.Z. designed the research. C.S. performed the research and analyzed the data. C.S. and T.Z. wrote the article.

## ACKNOWLEDGMENTS

This work was supported by grants 11931019 and 11775314 from Natural Science Foundation of P. R. China.

